# When does a parasite become a disease? eDNA unravels complex host-pathogen dynamics across environmental stress gradients in wild salmonid populations

**DOI:** 10.1101/2024.05.07.592920

**Authors:** Duval Eloïse, Blanchet Simon, Quéméré Erwan, Jacquin Lisa, Veyssière Charlotte, Loot Géraldine

## Abstract

Infectious diseases stem from disrupted interactions among hosts, parasites, and the environment. Both abiotic and biotic factors can influence infection outcomes by shaping the abundance of a parasite’s infective stages, as well as the host’s ability to fight infection. However, disentangling these mechanisms within natural ecosystems remains challenging. Here, combining environmental DNA analysis and niche modeling at a regional scale, we uncovered the biotic and abiotic drivers of a lethal infectious disease of salmonid fish, triggered by the parasite *Tetracapsuloides bryosalmonae.* We found that the occurrence and abundance of the parasite in the water—i.e., the propagule pressure— were mainly correlated to the abundances of its two primary hosts, the bryozoan *Fredericella sultana* and the fish *Salmo trutta*, but poorly to local abiotic environmental stressors. In contrast, the occurrence and abundance of parasites within fish hosts—i.e., proxies for disease emergence—were closely linked to environmental stressors (water temperature, agricultural activities, dams), and to a lesser extent to parasite propagule pressure. These results suggest that pathogen distribution alone cannot predict the risk of disease in wildlife, and that local anthropogenic stressors may play a pivotal role in disease emergence among wild host populations, likely by compromising the hosts’ ability to fight the parasite. Our study sheds light on the intricate interplay between biotic and abiotic factors in shaping pathogen distribution and raises concerns about the effects of global change on disease emergence.

## Introduction

The outcomes of host-parasite interactions strongly depend on the surrounding environmental conditions (Wolinska & King, 2009). In healthy ecosytems, host-pathogen dynamics often result in co-adaptation between the host and the pathogen, thus leading to limited negative impacts on host fitness. The host exhibits resistance and/or tolerance to the parasite, and the pathogen can persist in the environment without causing detrimental effects on host populations. However, rapid and drastic changes in environmental conditions due to human activities can affect key parameters such as parasite survival, virulence and transmission rate, as well as host resistance/tolerance to infection (Altizer et al., 2013; Budria & Candolin, 2014). In other words, environmental disturbances can disrupt “benign” host-parasite dynamics, favouring the emergence or the resurgence of infectious diseases with severe deleterious impacts on wild host populations (Schrag & Wiener, 1995; Lafferty, 2009; Gallana et al., 2013; Altizer et al., 2013). Understanding under which environmental conditions and through which mechanisms parasites impact their hosts and cause emerging diseases is thus critical to anticipate host health issues and demographic declines in animal and human populations.

Environmental stressors driving host-parasite dynamics include a range of abiotic factors acting on host behaviour or physiology, on the parasite inside its hosts (especially in ectotherms), and/or on the parasite outside its hosts during its free-living stages in the environment. The probability of host-parasite encounter and subsequent infection outcomes are also mediated by biotic factors such as the density of parasite propagules in the environment (hereafter, the propagule pressure) and the density of hosts in the environment (Pietrock & Marcogliese, 2003; Lootvoet et al., 2013; Lagrue & Poulin, 2015). For instance, the higher the parasite propagule pressure, the more likely the infection by the hosts. Reciprocally, the higher the host(s) density, the more efficient the parasite life-cycle (Arneberg et al., 1998; Hallett et al., 2012; Lootvoet et al., 2013). Acting synergistically, biotic and abiotic factors shape the parasite occurrence and abundance in both the environment and within its hosts, thereby driving the impact of parasites on host populations (Turner et al., 2021). A change in one or a few of these environmental factors may lead to increased infection rate and/or increased pathogenecity (Martin et al., 2010; Budria & Candolin, 2014; Cable et al., 2017). For instance, Johnson et al. (2007) identified cascading effects of water eutrophication on the outcomes of the trematode parasite *Ribeiroia ondatrae* infection in its amphibian host *Rana clamitans*. Eutrophication promoted algae development, increasing the density of snail intermediate hosts *Planorbella trivolvis*, which in turn produced and released more infective stages of the pathogen in the environment (higher propagule pressure), which ultimately increased infection intensity in the amphibian host population. Other studies found that increased water temperature negatively affected the immune capacity of amphibian hosts, which increased their susceptibility to infection by the deadly fungus *Batrachochytrium dendrobatidis* (Raffel et al., 2006; Rohr & Raffel, 2010). To understand when and how seemingly benign host-parasite interactions can cause large disease outbreaks in natural populations, it is thus important to disentangle the respective effects of the environmental factors (biotic and abiotic) on both parasite exposure and host susceptibility (James et al., 2015; Stewart Merrill et al., 2021).

Acquiring knowledge on the distribution (occurrence and abundance) of the parasite propagule pressure and its underlying environmental drivers is therefore one of the keys to understand and forecast disease outbreaks, especially for pathogens that are transmitted through the environment (Cable et al., 2017; Marcogliese, 2008; Okamura & Feist, 2011). However, one current technical limitation is that parasites are usually quantified within their host organisms (as prevalence or intensity estimates), but rarely as free-living stages in the environment. There is thus often a lack of information about the host exposure to infective propagules. The primary reason is that free-living stages are particularly challenging to detect due to their microscopic size and high dilution in the environment, which complicates their detection and quantification. The development of molecular detection techniques related to environmental DNA (eDNA) has revolutionised the biomonitoring and/or surveillance of rare and cryptic species, as well as the early detection of invasive species (Bohmann et al., 2014; Rees et al., 2014). Recent improvements in eDNA methods enable quantifying the abundance (or relative abundance) of a target species (Lodge et al., 2012; Doi et al., 2015; Seymour, 2019). Accordingly, eDNA has become an important tool in parasitology to improve the detection of otherwise invisible pathogens (Huver et al., 2015; Bass et al., 2015). For instance, Carraro et al. (2017, 2018) used eDNA to unravel patterns of occurrence of *Tetracapsuloides bryosalmonae*, an emerging myxozoan parasite of salmonid fish, in an alpine river. Detecting parasite DNA in the open water is thus a promising avenue to quantify the exposure of hosts to parasite propagules.

In this study, we investigated the mechanistic pathways explaining the emergence of the proliferative kidney disease (PKD) caused by the myxozoan parasite *T. bryosalmonae* in salmonids. This disease leads to massive mortality events worldwide both in aquaculture and in the wild (20-100% of mortality, Okamura et al., 2011). Our first objective was to identify the biotic and abiotic drivers of the parasite distribution in the environment (occurrence and abundance of parasite propagules, III and III, Fig. 1). The life cycle of *T. bryosalmonae* involves two successive hosts: a salmonid fish and a bryozoan. The parasite has been found in bryozoans even in the absence of intermediate fish hosts. Its final bryozoan hosts therefore represent a pervasive reservoir for future fish infection (Okamura et al., 2001). As with most parasites, we expected that the distribution (occurrence and abundance) of *T. bryosalmonae* in the water, as estimated from eDNA would be strongly influenced by the local abundances of its two hosts (the fish *Salmo trutta* and the bryozoan *Fredericella sultana;* III, Fig. 1). We also expected water temperature, which determines the amount of spore released from the bryozoan, to affect parasite distribution (Wahli et al., 2008; III, Fig. 1). Our second objective was to characterize the biotic and abiotic factors determining parasite infection within host populations (@ and O, Fig. 1, occurrence and abundance in individual fish hosts). Assuming that the parasite DNA abundance measured in the environment is a reliable proxy for the parasite propagule pressure, we tested the relative role of the parasite propagule pressure and the most prevalent abiotic stress factors, on the parasite occurrence and abundance in the fish host (as indicators of disease development). We expected abiotic stressors (such as high water temperature) to trigger disease development in fish either by favoring the abundance of free-living infective stages in the water (indirect impact of the abiotic environmental stressors on propagule pressure, III and O, Fig 1), and/or by altering the immune and physiological ability of the fish host to resist/tolerate infection (direct impact of the abiotic stressors on fish resistance/tolerance, @, Fig. 1). To test these predictions, we combined eDNA methods and niche modelling across multiple sites and at a large geographic scale, and we compared niche models including the biotic, abiotic factors or both to assess their relative importance in explaining infection in wild fish populations. This original and integrative study, by harnessing the power of eDNA detection and niche models accounting both for the presence/absence and abundance of DNA in samples (Martin et al., 2005), helped disentangling and anticipating the effects of multiple environmental stressors on disease emergence in wild populations.

**Figure 1.**
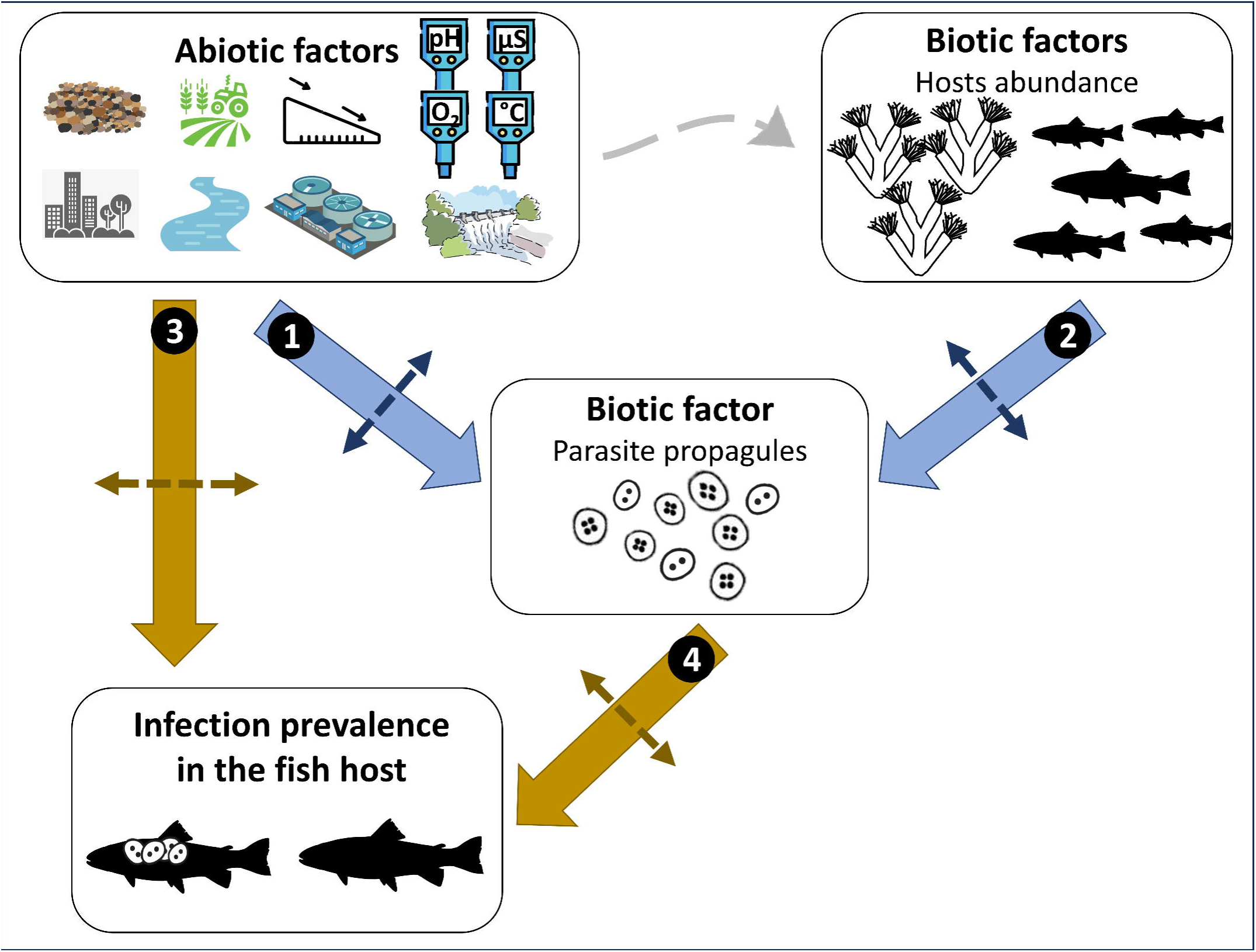
Visual representation of our research questions. First (IL, IL), we investigated the abiotic (environmental factors) and biotic factors (hosts abundance) responsible for *T. bryosalmonae*’s distribution (occurrence and abundance) in the environment (parasite propagule pressure). Second, (@, O), we investigated the abiotic (environmental factors) and biotic (parasite propagule abundance) factors responsible for the distribution (occurrence and abundance) of the *T. bryosalmonae* in brown trout host (infection patterns and disease emergence). Abiotic factors may act directly on infection within fish by acting on fish physiology, but also indirectly by determining the parasite propagule abundance.

## Methods

### Host-parasite system

*T. bryosalmonae* needs two hosts to complete its life cycle: a bryozoan (definitive host, here *Fredericella sultana*, its main and most widespread bryozoan host in our study area, Schmidt-Posthaus et al., 2021, Hartikainen, pers. comm.) and a salmonid fish (intermediate host, here *S. trutta*, its only fish host in the area) (Okamura et al., 2011). The transitions between its life stages are temperature-dependent. Parasite propagules release in the river by the bryozoans occurs when the water temperature reaches 9°C (Gay et al., 2001), with peaks in spring and autumn (Tops et al., 2009; Duval, 2022). The released parasite propagules infect brown trout by entering through gills and skin, circulate through the blood until reaching the kidney where they develop, potentially triggering an exaggerated immune reaction of the fish host when water temperature exceeds 15°C, leading to PKD development, especially during summer (Hedrick et al., 1993). The disease may develop at the first infection of naive fish, and if they survive, they acquire immunity upon reinfection, so that young-of-the-year fish are the most sensitive stage (Feist & Longshaw, 2006). The severity of PKD following *T. bryosalmonae* infection in brown trout may also depend on water temperature, and the disease in turn modulates brown trout thermal tolerance and metabolic rate because of decreased erythropoiesis leading to anemia (Okamura et al., 2011; Bruneaux et al., 2017). In addition, brown trout is a cold-water species so that increasing temperature can also trigger physiological stress (Elliott & Elliott, 2010). Water quality is also an important environmental parameter potentially affecting host-pathogen interactions. Indeed, the development of the parasite and its bryozoan host is favored by the quantity of nutrient available in the stream, while brown trout physiology is negatively affected by increased nutrients and decreased oxygen rate (Hartikainen et al., 2009; Bailey et al., 2018; Rubin et al., 2019; Duval, 2022). This suggests complex interplays between biotic and abiotic environmental factors on PKD disease dynamics.

### eDNA sampling

The study covered a wide area in Southern France, with 83 sites scattered along an East-West gradient in the Pyrenean Mountains and over 54 streams (Fig. 2). We covered an altitudinal band from 230 to 940m a.s.l., corresponding to the altitudinal range in which the parasite is generally found (it is rarely detected at altitudes >800m). We sampled eDNA between the 30^th^ of July and the 14^th^ of August 2020 to estimate the abundances of the two hosts (*S. trutta* and *F. sultana*) and of the parasite in the environment during the most favourable period for PKD development.

**Figure 2.**
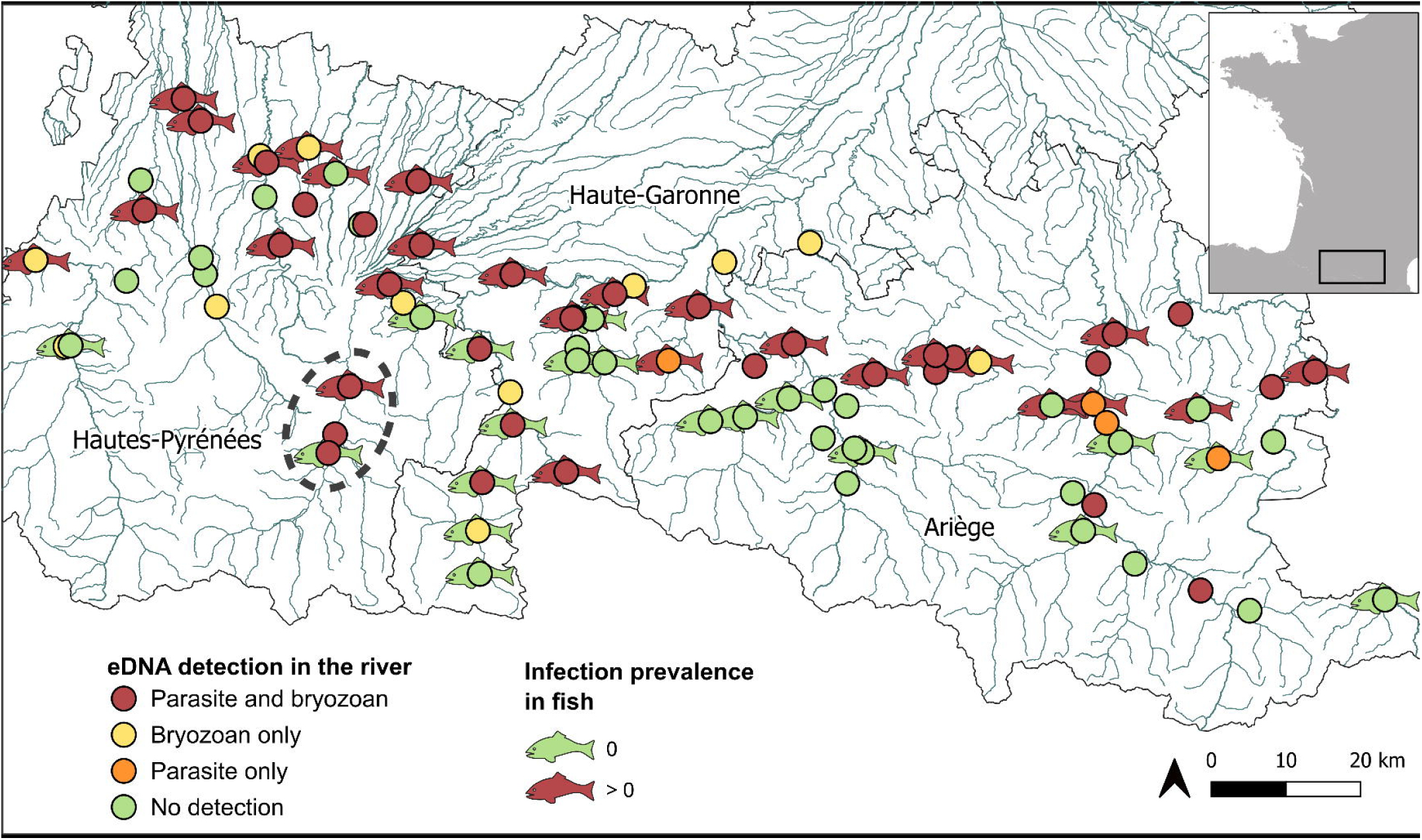
Map of the 83 sampled sites. The presence or absence of detection of the parasite (*T. bryosalmonae*) and its bryozoan host (*F. sultana*) in the water is represented by circles, and its presence/absence in the fish host is indicated by fishes. Inset indicates the location of the studied area, at the South of France. The dotted ellipse examplifies three sites sampled on the Neste river described in the discussion.

At each site, we filtered up to 12L of water onto 1.2µm cellulose nitrate Sartorius® filters (Ø 50mm) with a Vampire sampler (Bürkle®) and Sartorius® filter holders as follows: we used 8 filters per site and filtered a maximum of 1.5L per filter, less when filter clogging prevented it, in which cases we measured the volume of water filtered per filter. We stored pairs of filters in 5mL Eppendorf® tubes, to get 4 field replicates per site that were quickly stored at -80°C until DNA extraction.

### DNA extraction and amplification

To measure the occurrence and abundance of the pathogen spores in the water, as well as the abundance of the intermediate (trout) and final (bryozoan) hosts in the environment, we used multiplex droplet digital PCR (ddPCR) assays for the detection of the three species from water eDNA. We performed DNA extraction directly on filters using QIAGEN® PowerSoil kit following manufacturer recommendations and under strict laboratory environment required for eDNA extractions. We used the primers and probes designed by Carraro et al. (2018) and Carim et al. (2016) to amplify a 71bp fragment of *F. sultana* 16S SSU rDNA sequence, a 102bp fragment of *T. bryosalmonae* COI DNA and a 108 bp fragment of *S. trutta* cytochrome B DNA (Table 1). Target DNA was amplified using a BioRad QX200 Droplet Digital PCR system™ (Bio-Rad, Temse, Belgium), with the following thermal conditions: 10min at 95°C, then 40 cycles encompassing 30s at 94°C and 1min at 60°C, followed by 10min at 98°C and 30min at 4°C. The PCR reactions were performed on a total volume of 22µL including 11µL of EvaGreen digital PCR Supermix, 2.4µL of sample DNA and 8.6µL of primer mix (including 1.9µL of each primer and 0.5µL of each probe, 10µM). Each 96- well run included 4 PCR negative controls with water only, and 1 PCR positive control consisting of *F. sultana* tissue infected by *T. bryosalmonae*. The baseline threshold for separating positive and negative droplets was manually chosen for each ddPCR run, according to the distribution of the droplets from the negative and positive control wells. We run 2 ddPCRs per sample: one with the primers and probes amplifying *F. sultana* and *T. bryosalmonae* and one with the primers and probes amplifying *S. trutta* and *T. bryosalmonae*. We targeted *T. bryosalmonae* DNA twice to maximise the chances of detection of this species for which we expected low concentrations in the water (Sieber et al., 2020).

**Table 1.**
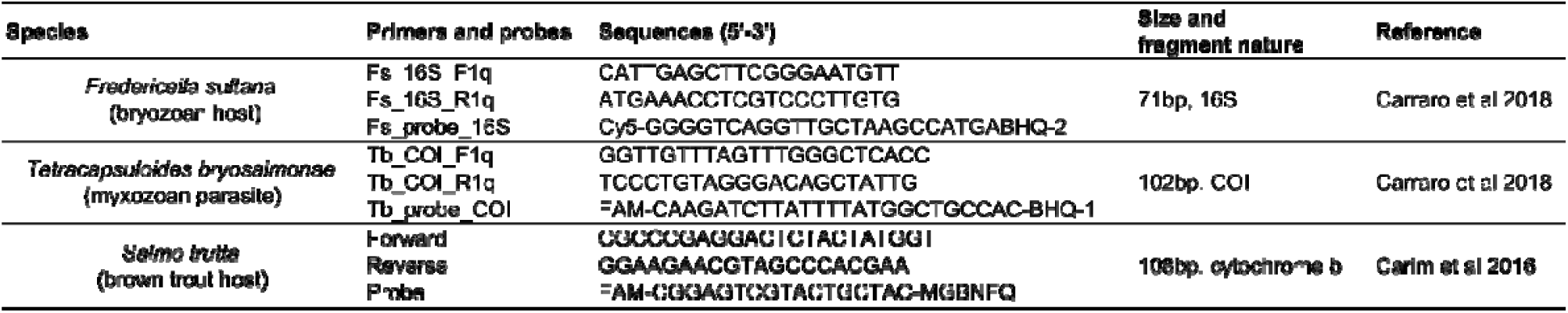
Primers and probes sequences used to amplify *F. sultana*, *T. bryosalmonae* and *S. trutta* DNA, either in water samples or urine samples from the fish.

### Infection prevalence and abundance

To measure *T. bryosalmonae* occurrence and abundance in juvenile brown trout populations, we sampled fish at 46 of the 83 sites sampled for eDNA. We could not cover all the sites because electro-fishing is time-consuming and mobilizes a substantial team on the field. For each of these 46 sites, we sampled up to 20 individuals (mean±SD, 18±3 individuals). We targeted juvenile trout (mean size±SD, 78±16mm), corresponding mainly to young-of-the-year (0+) fish because it is the most abundant and sensitive stage. We used a non-lethal method (uDNA for urine DNA) based on the excretion of *T. bryosalmonae* spores by infected fish through urine excretion to infer the infection status of each fish (whether spores were released or not) and the severity of the infection of each fish (assuming that a higher abundance of spores released by the fish corresponds to a more severe infection). Details about the uDNA method are available from previous studies (Duval et al., 2021, 2022). Here, and for later analyses, we focus only on the mean abundance of spores released by fish averaged at each site. We also ran models using the infection prevalence of fish (number of infected fish divided by total number of sampled fish per site) as a response variable but they are not presented here (as they yielded similar conclusions). All fish were then released alive into their sites of sampling. Authorisations to sample brown trout were provided by the Directions Départementales des Territoires of Ariège, Haute-Garonne and Hautes-Pyrénées, respectively.

### Environmental data for niche modelling

A wide range of environmental factors were measured or extracted from available databases for each sampling site to assess their impact on *T. bryosalmonae* abundance, in the environment and in the fish host. We used an In-Situ® Aqua TROLL 500 Multiparameter Probe to measure water temperature, pH, specific conductivity and O_2_ concentration at each site during the eDNA survey (summer 2020). We used QGIS software (2022) to get information on the land use with the CORINE Land Cover 2018 dataset (European Environment Agency) on a 2km buffer around each site and we collected the percentage of forest, urban and agricultural land, as land use may impact water quality through nutrient input and chemical pollution (Tong & Chen, 2002). We used the Réseau Hydrographique Théorique (RHT, Pella et al., 2012) to get information on the mean flow (module), the sediment fineness (the higher the value, the finer the sediment), the river width and depth. The mean slope (‰) of the upstream 2kms was computed. As proxies for the impacts of human activities, we included the presence of dams, calculated as the cumulative height of dams and weirs 2kms upstream of the sampled site (hereafter “cumulative dams height”), as well as the cumulative nominal capacity of the Wastewater Treatment Plants (WWTP) 2kms upstream of the sampled sites (using the SANDRE Service d’Administration Nationale des Données et Référentiels sur l’Eau database) (Carey & Migliaccio, 2009; Zaidel et al., 2021). The mean flow, the cumulative dams height and the cumulative capacity of WWTP were log-transformed to homogenise their distribution.

We checked for correlation between the environmental variables and removed those that had a correlation coefficient>|0.7| to limit collinearity issues in subsequent models. We thereafter removed from the dataset the percentage of forested area (keeping the percentage of agricultural area that was inversely correlated) and the river width and depth. These two latter variables were strongly correlated with -and thus represented by- the mean water flow.

### Statistical analyses

All statistical analyses were conducted in the R environment (R 4.0.3; R Core Team 2020).

We divided the raw eDNA concentration of the three species by the number of liters filtered on the field, and averaged the concentrations across the four field replicates. The eDNA concentrations were multiplied by 100 and 10000 for *S. trutta* and *T. bryosalmonae*/*F. sultana* respectively to transform concentrations into count data and ease modelling. The DNA concentration of each species was log-transformed when used as an explanatory variable.

To test the determinants of *T. bryosalmonae* occurrence and abundance in the water (Ill and Ill, Fig. 1), we used hurdle linear models with negative binomial distribution (Hu et al., 2011; Loeys et al., 2012) to account for the excess of zeros in the distribution of *T. bryosalmonae* eDNA concentration. These models relate the eDNA concentration (as count data) and the environmental variables in two parts: a binary part modelling the occurrence (i.e., presence/absence) of the parasite, and a negative-binomial part modelling the abundance when the parasite is present. Although still underused in the context of eDNA data, this type of modeling approach appears particularly suited for species with sparce distribution, which is often the case of parasites, species with stringent environmental requirements or rare species (Potts & Elith, 2006). Here, environmental predictors include the DNA abundance of the two hosts (biotic factors), as well as the abiotic factors listed above. After visual exploration of the dataset, we included a polynomial term for the effect of water temperature in the models, as we identified a potential non-linear relationship with *T. bryosalmonae* abundance in the water.

We used a model selection procedure based on the small-sample size corrected Akaike Information Criterion (*AICc*, Burnham & Anderson, 2002) with the *MuMIn* package (Barton, 2020) to identify the most relevant variables sustaining the *T. bryosalmonae* eDNA distribution in the environment. During this process, we limited the selection procedure to models including no more than eight parameters (excluding the intercept) to avoid over-parametrization. We kept models with Δ*AICc*<4 relative to the best model and computed the relative importance (*RI*) of each variable as the cumulative weight of each model in which it appears (Burnham & Anderson, 2002). The cumulative weight of the model selection with Δ*AICc*<4 was standardized so that the *RI* of each term varied between 0 and 1. We considered that a variable was biologically-relevant when *RI*>0.5 (De Kort et al., 2021). We then used a model averaging procedure to compute the mean estimate of each relevant variable, averaging the estimates of the models in which it appeared (i.e., with the subset method).

To investigate the determinants of *T. bryosalmonae* occurrence and abundance in fish hosts (@ and O, Fig. 1), we used the same model selection procedure (and same type of models) as above to relate abiotic predictors and the abundance of parasite DNA in the water (a proxy for the parasite propagule pressure, biotic predictor) to the occurrence and abundance within the fish. Given that the dataset was more restrained, we limited the selection procedure to models including no more than seven parameters (excluding the intercept) to avoid over-parametrization.

To further assess the relative roles of the abiotic and biotic factors in determining the *T. bryosalmonae* occurrence and abundance measured in the environment and in the fish hosts respectively, we compared the predictive power (r^2^) between models including either only abiotic environmental variables or only biotic variables (among the variables with a *RI*>0.5), and compared their respective fit to the data using likelihood ratio tests.

## Results

### General patterns

As expected, *S. trutta* eDNA was detected at all sampling sites, confirming the presence of fish hosts in all sampling sites, and concurrently validating the reliability of our eDNA sampling and conservation protocols. The mapping of the occurrence of *T. bryosalmonae* and the bryozoan *F. sultana* in the water revealed that most of the time (96.2%), *T. bryosalmonae* was detected together with its bryozoan host (red dots, Fig. 2), except for 4 out of the 83 sites where it was detected alone (orange dots, Fig. 2). More than half of the trout populations (27 out of the 46 sites) were infected with *T. bryosalmonae.* Co-occurrences between *T. bryosalmonae* in the water and in the fish host were observed at 21 out of the 27 sites with infected fish (77%). In some cases, the parasite was detected in the water but not in the fish host (5 out of the 19 sites without infected fish, 26%, Fig. 2), and, more surprisingly, in some cases the parasite was detected in the fish host but not in the water (6 out of the 27 sites, 22%, Fig. 2. We detected neither the bryozoan nor the parasite in 29 out of the 83 sampled sites (Fig. 2).

### Parasite DNA occurrence and abundance in the water (proxy for parasite propagule pressure)

The model selection revealed that the most likely variables to explain the occurrence and abundance of *T. bryosalmonae* in the water are the abundance of the two hosts (biotic factors), water conductivity, and the cumulative height of dams (abiotic factors) (all *RI>*0.75, Fig. 3a). The DNA abundances of trout and bryozoan in the water showed the highest *RI* in explaining both the occurrence and abundance of *T. bryosalmonae* (Fig. 3a), emphasizing the importance of biotic factors for pathogen distribution in the environment. More specifically, the occurrence of *T. bryosalmonae* at a site increased with the DNA abundance of the two host species, and once settled at a site, *T. bryosalmonae* abundance also increased with the DNA abundance of the two hosts (Fig. 3b).

**Figure 3.**
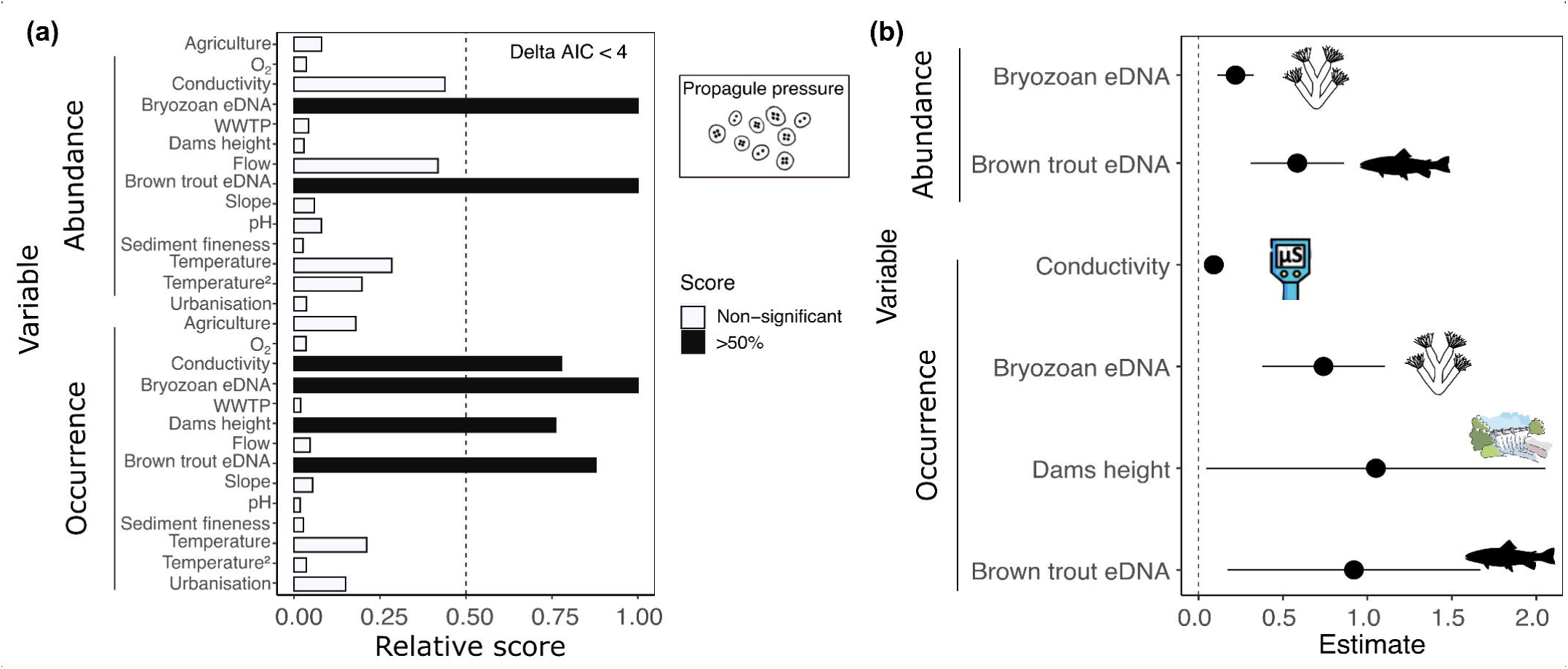
**(a)** Relative importance (*RI*) of environmental factors investigated to explain the occurrence and abundance of *T. bryosalmonae* in the water. The relative importance of each factor is estimated as the standardized (between 0 and 1) cumulative weight of each model in which a given factor appears. The dashed grey line indicates the 0.5 threshold. **(b)** Mean estimates of the effects of each relevant variable (*RI* > 0.5) on *T. bryosalmonae* abundance and occurrence in the water. The 95% confidence intervals are indicated. The dashed black line represents a null effect.

In addition, the occurrence of *T. bryosalmonae* at a site also tended to be higher when the water conductivity was high and when there was a significant presence of dams 2km upstream (Fig. 3b). However, the comparison of models including either abiotic or biotic factors alone revealed that the model including only the two host DNA abundance variables explained much more variance in the distribution of *T. bryosalmonae* in the water than the model including only the abiotic factors (51% vs. 5% respectively, χ*²=*69.11, *df=*2, *P*<0.001). This indicates that the occurrence and abundance of *T. bryosalmonae* DNA in the water was mainly driven by the abundances of its two hosts in the environment, and poorly by the surrounding environmental conditions.

### T. bryosalmonae infection in fish (proxy for disease emergence)

The model selection procedure revealed that abiotic environmental factors such as sediment fineness, water temperature, percentage of agricultural lands and cumulative height of dams and the abundance of *T. bryosalmonae* spores in the water, are the most likely variables explaining the occurrence and abundance of *T. bryosalmonae* in the fish host (RI>0.5, Fig. 4a). More specifically, the occurrence of *T. bryosalmonae* in fish host increased in sites with higher agricultural activities and finer sediments, and in sites with higher abundance of *T. bryosalmonae* in the water (Fig. 4b). Once settled in fish populations, the abundance of *T. bryosalmonae* in fish was higher in warmer sites, in sites with higher abundance of *T. bryosalmonae* in the water, and in sites with a lower height of dams upstream (Fig. 4b).

**Figure 4.**
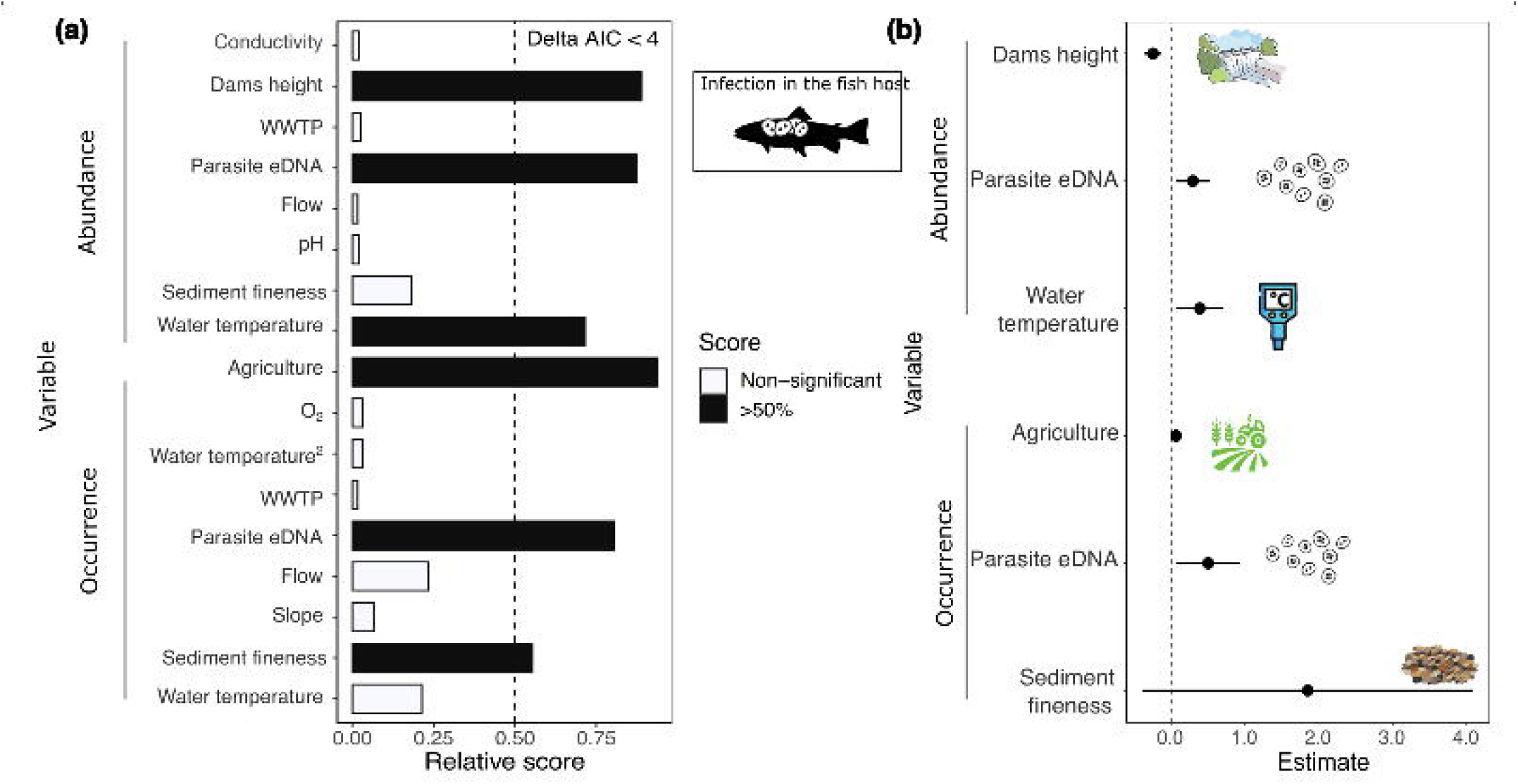
**(a)** Relative importance (*RI*) of environmental factors investigated to explain the occurrence and abundance of *T. bryosalmonae* in the fish host (*S. trutta*). The relative importance of each factor is estimated as the standardized (between 0 and 1) cumulative weight of each model in which a given factor appears. The dashed grey line indicates the 0.5 threshold. **(b)** Mean estimates of the effects of each relevant variable (*RI* > 0.5) on the abundance and occurrence of *T. bryosalmonae* in the fish host. The 95% confidence intervals are indicated. The dashed black line represents a null effect.

The comparison of models including either abiotic or biotic factors alone revealed that the model including only the abiotic factors explained twice as much of the variance as the biotic-model, which solely included the abundance of *T. bryosalmonae* in the water (47% vs. 23% respectively, χ*²=*31.86, *df=*2, *P*<0.001). This shows that *T. bryosalmonae* infection in brown trout was primarily influenced by abiotic environmental factors acting directly on the hosts, especially water temperature and agricultural activities, while the impact of the parasite propagule pressure in the water was relatively lower.

## Discussion

We have developed an innovative methodological framework that combines eDNA methods and large-scale environmental niche modelling accounting for both the occurrence and abundance of key species to explore the abiotic and biotic factors underlying disease emergence in aquatic wildlife. This integrative framework enables us to encompass all mechanistic pathways from the distribution of the parasite in the environment to fish infection. Our results pointed out that the abundances of the two main hosts were the most important factors driving the occurrence and abundance of *T. bryosalmonae* propagules in the water. In contrast, *T. bryosalmonae* infection within the brown trout host was strongly driven by abiotic factors such as temperature and agricultural activities. Our results imply that high abundances of parasite propagules pressure in the environment are not solely responsible for disease emergence, and that abiotic stressors linked to human activities play a pivotal role in disease emergence in the wild, likely by influencing host health and resistance and/tolerance to the pathogen.

### Pathogen distribution in the water

Our findings demonstrated that water eDNA is a particularly valuable tool for large-scale spatial surveillance of free-living forms of pathogens in the environment, which are often difficult to detect using conventional approaches. It is also an unparalleled approach for understanding the factors driving the co-occurrence of parasites and hosts along gradients of environmental stress. A major finding of our study is that both the occurrence and abundance of *T. bryosalmonae* DNA in the water (a proxy for the parasite propagule pressure) were strongly and positively associated with the abundances of its bryozoan and fish hosts. Assuming that DNA concentrations found in the water are a good proxy for species abundances, as confirmed by previous eDNA studies using species-specific markers (Yates et al. 2019), this strongly suggests that higher abundances of *F. sultana* and *S. trutta* correlate with an increased likelihood of *T. bryosalmonae* colonization at a site, leading to higher abundance of *T. bryosalmonae* in the water once settled. We anticipated this strong association with bryozoan abundance because it is *T. bryosalmonae*’s definitive host (Okamura et al., 2011). However, the strong association with brown trout abundance was rather unexpected. Previous studies suggested that the parasite DNA detected in the water may primarily originate from bryozoan release (Carraro et al., 2017, 2018), but these studies did not estimate brown trout abundance. This suggests that fish host could also contribute to amplifying the abundance of pathogen spores in the water, probably through important spore release in urine after amplification within fish kidneys (parasite target organ). Further investigations to determine the exact nature of pathogen spores found in the water (infectious spores released by bryozoans, or spores released by the fish) are in progress to address this question.

In addition, the occurrence of *T. bryosalmonae* DNA in the water was positively associated with the presence of dams upstream and water conductivity. These environmental conditions may be particularly suitable for the bryozoan growth and for parasite release, and may hence boost locally the colonization by *T. bryosalmonae*. Indeed, dams may favor bryozoan colonies (due to the lentic nature of the habitat) and may warm-up the water locally which supposedly favors the bryozoan and parasite life cycles. Similarly, a high water conductivity is generally associated with high nutrient loads, which may also be favorable for bryozoan colonies (Hartikainen et al., 2009; Ros et al., 2022). Nonetheless, the presence of the two hosts (55% of the total variance of the occurrence and abundance of *T. bryosalmonae*) largely outweighed abiotic factors (6%) in shaping the distribution of *T. bryosalmonae* in the water, which supports the idea that most parasites rely primarily on the presence of hosts for survival and reproduction (Staniczenko et al., 2017; Facon et al., 2021).

### Determinants of pathogen infection in fish

In our study area, heavily infected fish populations with high parasite prevalence (>90%) and load typically develop major pathological lesions and experience increased mortality rate (Garmendia & Lautraite, 2017). Parasite occurrence and abundance within fish measured through our non-lethal uDNA approach are hence a good proxy for the emergence of the disease. Our results suggest that the occurrence and abundance of *T. bryosalmonae* in fish host populations and therefore, the epidemiological dynamics of the PKD disease at the regional scale, were mostly driven by abiotic environmental conditions (47% of the total variance explained), although we also revealed a positive -but surprisingly weaker- influence of the abundance of parasite propagules in the water (23% of the total variance explained). This corroborates a recent experiment revealing that fish parasite load (measured in the kidney) did not differ between fish groups exposed either to low or high parasite spore concentrations (Strepparava et al., 2020). In some upstream sites, host-parasite system seems “balanced”, with the parasite and its hosts coexisting, but with no or very few trout infected (Fig. 2). Conversely, our results suggest that in downstream sites, alterations of abiotic conditions could disrupt this balance and favor the emergence of the disease. Indeed, we found that both the percentage of agricultural landscape and increased water temperature positively correlate with *T. bryosalmonae* infection in fish (occurrence and abundance in the urine respectively). These findings likely reflect impaired fish physiology and immunology under stressful conditions, which indirectly increases parasite proliferation within the fish host (Bruneaux et al., 2017; Lauringson et al., 2021). This is consistent with previous studies indicating that agricultural pollution (*sensu lato*) and water temperature are major environmental stressors for brown trout, with negative consequences for immunological, metabolic and physiological defense parameters and hence for their ability to resist pathogens (Bruneaux et al., 2017; Bailey et al., 2017; Borgwardt et al., 2020; Waldner et al., 2021).

In addition to these two stressful factors (agriculture and temperature) that likely alter fish defenses, we found that fish infection tended to increase in sites with finer sediments. This is somewhat consistent with a previous study in alpine streams showing a strong association between bryozoan development and the substrat type (Carraro et al., 2018), and suggests that physical characteristics of the riverbed might partly control host-parasite dynamics. We also observed that the presence of dams decreased the abundance of spores released by the fish, which is surprising given that dams increased the occurrence of *T. bryosalmonae* spores in the water (as discussed earlier). One could hypothesize that sediment size and dams have complex indirect effects on disease dynamics, for instance by favoring the development of bryozoans and/or the contact rate between parasite spores and fish hosts. Accordingly, Mathieu-Bégné et al. (2021) experimentally found a strong influence of the substrate composition at the microhabitat scale on the infection of the rostrum dace *Leuciscus burdigalensis* by the crustacean ectoparasite *Tracheliastes polycopus*, defining what they called “hotspots of infection”. However, given the correlative nature of our study, these findings must be interpreted with caution. Further local-scale and/or experimental approaches are now needed to refine these findings and reveal underlying mechanisms of such infection hotspots. Importantly, all these abiotic effects were partly independent from the parasite propagule pressure, which suggests that measuring parasite DNA concentration in the water is not sufficient to inform on the health status of the host populations, but rather informs on the risk of disease emergence under adverse environmental conditions.

### Potential applications for PKD outbreak surveillance

Beyond the underlying mechanisms, a striking result from this large-scale survey is that the emergence of PKD in brown trout populations is not solely driven by the abundance of parasites in the water. Indeed, even a low abundance of parasite in the water can lead to strong disease risk if abiotic conditions are unfavorable for the fish host. This raises interesting avenues for conservation actions to limit disease risk and population collapse in salmonid populations exposed to PKD in Europe. For instance, focusing on water quality and limiting nutrient input and temperature increase could help improving fish defense to the disease while acting on sediment and/or dams could limit the presence of bryozoans and/or pathogen contact rate.

In addition, the mapping of *T. bryosalmonae* and its bryozoan host in the water reveals sites where future PKD outbreaks might occur in fish, *i.e.*, sites in which either the bryozoan, the parasite or both are detected in the water but for which infection in fish is not detected yet. For instance, in the Neste River (the sites included in the dotted ellipse in Fig. 2), parasite eDNA is detected at all the three sampling sites, whereas there is no fish infection at the uppermost site (which has been confirmed by independent measures of infection directly in the kidney, A. Lautraite, pers. comm.). These sites should be prioritized for pro-active surveillance to avoid future outbreaks, demonstrating the importance of eDNA as an operational tool for environmental managers.

### Conclusions

We demonstrated here the usefulness of eDNA (and uDNA) to (i) map the large-scale distribution of an emerging fish pathogen in both the water and the vertebrate host and thus ultimately map disease risk in wildlife (ii) reveals the abiotic and biotic drivers and processes making this pathogen a harmful disease for brown trout populations. By quantifying simultaneously the different forms *T. bryosalmonae* pathogen (in the water as propagule pressure and in the fish host), we further showed that the presence of parasite in the water is not sufficient to predict infection in fish. Indeed, our results suggest that disease risk is triggered by particular environmental conditions altering host physiology, and/or parasite multiplication inside the host, and/or the contact rate between infective stages and the hosts. Using non-lethal approaches, this integrative and large-scale study reveals the importance of biotic factors (host abundance) for the parasite life-cycle, and the importance of abiotic conditions (environmental stressors and riverbed characteristics) in shaping, directly and indirectly, the dynamics of an emerging infectious disease in the wild. In addition, this will hopefully help building operational tools for biodiversity managers to limit emerging disease risk under global change.

